# Effects of more natural housing conditions on the muscular and skeletal characteristics of female C57BL/6J mice

**DOI:** 10.1101/2022.09.27.509671

**Authors:** Paul Mieske, Julia Scheinpflug, Timur Alexander Yorgan, Laura Brylka, Rupert Palme, Ute Hobbiesiefken, Juliane Preikschat, Kai Diederich, Lars Lewejohann

## Abstract

**Background:** Enrichment of home cages in laboratory experiments offers clear advantages, but has been criticized in some respects. First, there is a lack of definition, which makes methodological uniformity difficult. Second, there is concern that the enrichment of home cages may increase the variance of results in experiments. Here, the influence of more natural housing conditions on physiological parameters of female C57BL/6J mice was investigated from an animal welfare point of view. For this purpose, the animals were kept in three different housing conditions: conventional cage housing, enriched housing and the semi-naturalistic environment. The focus was on musculoskeletal changes after long-term environmental enrichment.

**Results:** The housing conditions had a long-term effect on the body weight of the test animals. The more complex and natural the home cage, the heavier the animals. This was associated with increased adipose deposits in the animals. There were no significant changes in muscle and bone characteristics except for single clues (femur diameter, bone resorption marker CTX-1). Additionally, the animals in the semi naturalistic environment (SNE) were found to have the fewest bone anomalies. Housing in the SNE appears to have the least effect on stress hormone concentrations. The lowest oxygen uptake was observed in enriched cage housing.

**Conclusions:** Despite increasing values, observed body weights were in the normal and strain-typical range. Overall, musculoskeletal parameters were slightly improved and age-related effects appear to have been attenuated. The variances in the results were not increased by more natural housing. This confirms the suitability of the applied housing conditions to ensure and increase animal welfare in laboratory experiments.

## 1. Background

Enrichment of the housing conditions of laboratory animals is not subject to any universal principles and therefore takes different forms (1). In general, increased provision of stimuli to enable natural behavior and improve animal welfare could be a definition of environmental enrichment (EE). In fact, many comparative studies on the impact of EE on laboratory mice show effects on animal welfare (2). The majority of studies conclude that the use of EE is beneficial for the animals. Housing in enriched environments can reduce stereotypies (3–7), reduce anxiety (8–10) and promote exploratory behavior (11, 12), as well as promote the development of favorable physiological parameters, especially in the neurological field (13, 14). It is also important that the enrichment is versatile and easily accessible. Dispersed enrichment using different elements or larger enclosures has the potential to minimize aggression between individuals in a group (15). Despite the clear advantages of using EE, the minimum legal requirements for standard laboratory husbandry unfortunately remain unchanged. In most experimental studies the cages continue to be only minimally equipped and thus constitute a barren environment. A recent meta-analysis has shown that such barren housing conditions in biomedical translational studies can negatively affect a number of health parameters in the experimental animals, thus substantially compromising the validity of these studies (16).

However, mandating enrichment of laboratory animal cages it is not an easy endeavor, as a number of factors must be considered for implementation. Laboratories and breeding facilities need to gauge which EE elements can be added, minding the applicability and financial feasibility. When introducing EE elements, hygiene standards for the animals must be maintained and standardization between experiments and laboratories must not be compromised (17). Finally, an increasing individualization of the cage design according to one’s own views, experiences, and tastes may affect the comparability of methods and results.

Another major criticism on the use of EE is the fear that the emergence of individual differences of experimental animals promoted by EE results in increased variability. However, previous studies have shown that the use of EE emphasizes individuality without necessarily increasing the variability of physiological parameters (18, 19). Providing simple enrichments such as additional nesting material and plastic or cardboard tunnels over long periods can change social behavior, but do not necessarily affect physiological parameters (20). A further advance to enriching conventional housing conditions of laboratory animals is to design an environment that resembles nature. However, even the use of a semi-natural environment that promotes individual diversified behavior did not result in greater deviations in the measurement of physiological parameters compared to conventional studies (21, 22).

Only by gaining more knowledge and disclosing additional benefits of EE is it possible to increase awareness and acceptance of the need to use EE, which ultimately also leads to an increase in overall animal welfare.

An example of a suitable method of laboratory cage enrichment was recently presented by Hobbiesiefken *et al.* (3). By exchanging enrichment elements in different categories on a weekly basis, an EE concept was developed, that reduced stereotypies and served to evaluate individual enrichment elements through behavioral observations. In the present study, three housing conditions with increasing opportunities to exhibit more natural behavior were used to analyze effects on physiological parameters. In addition to the conventional and enriched housing conditions used by Hobbiesiefken *et al.* a semi-natural environment (21–27) was used to exploit the full potential of the mice’s natural behavior as much as possible. Since many previous studies focus more on animal behavior than on uncovering the effects of EE on physiological traits, we here analyze whether the use of objects, social enrichment, or larger enclosure space affects musculoskeletal characteristics of female C57BL/6J mice. Nevertheless, our analysis of musculoskeletal characteristics was conducted within the framework of the animal welfare perspective. Additionally, to be able to monitor changes in animal welfare on a more obvious and holistic level, body weight, resting metabolic rate and stress hormone levels were measured. With regard to biomedical studies it might be under consideration, whether an increase or decrease of the respective parameter is the desired outcome. We admit that it might be questionable whether an altered muscle weight or bone density is ultimately decisive for improved animal welfare. However, one could assume that the physiological parameters we measured under more challenging environmental conditions are a better representation of the natural state. When considering effects on human bone structure, twin-studies are used to distinguish between genetic and environmental influences. It has already been established that in humans up to 70% of individual differences are of genetic origin (28, 29). In laboratory mice, genetic variability can be controlled. Thus, to pay greater attention to the influence of environmental enrichment on muscle and skeletal properties, inbred mouse strains are particularly suitable, since virtually an unlimited number of genetically identical individuals are available. Nonetheless, it has been shown that individual differences emerge despite genetic uniformity even in strictly standardized and limited housing conditions (30). Albeit genetics and housing conditions are not exclusive factors that should be considered for the evaluation.

## 2. Results

### 2.1. Weight data

On arrival, the experimental animals weighed 19.8 ± 0.9 g (4.5 %, 22.5 – 17.2 g, n = 44). During the experimental period of 88 weeks animal weight increased significantly (F(1|3604) = 7716, p-value < 0.001, R² = 0.68). A mixed model applied to the data confirmed that weight increased over time and is influenced by the individual animal, the housing condition and the time as a random effect and by the individual animal within the different housing condition and the individual animal over time as nested random effects. The applied model explains the variance of the data significantly better than a model that ignores the housing condition and time as predictors (p < 0.001). The different influences on the weight resulted in animals living in the SNE being the heaviest and the animals living in the CON being the lightest (Figure 1 A). At the time of perfusion of the animals they showed a mean animal weight in CON housing of 30.4 ± 3.1 g (10.1 %, 36.1 – 26.1 g, n = 11), in ENR housing of 33.5 ± 3.1 g (9.2 %, 40.5 – 29.8 g, n = 11) and in SNE housing of 34.6 ± 3.1 g (8.9 %, 40.0 – 28.4 g, n = 18, Figure 1 B). Animals in SNE housing were significantly heavier than the animals in CON housing. The weight increase from CON to ENR housing was marginally not significant. There was no significant difference between ENR and SNE housing.

**Figure 1.**
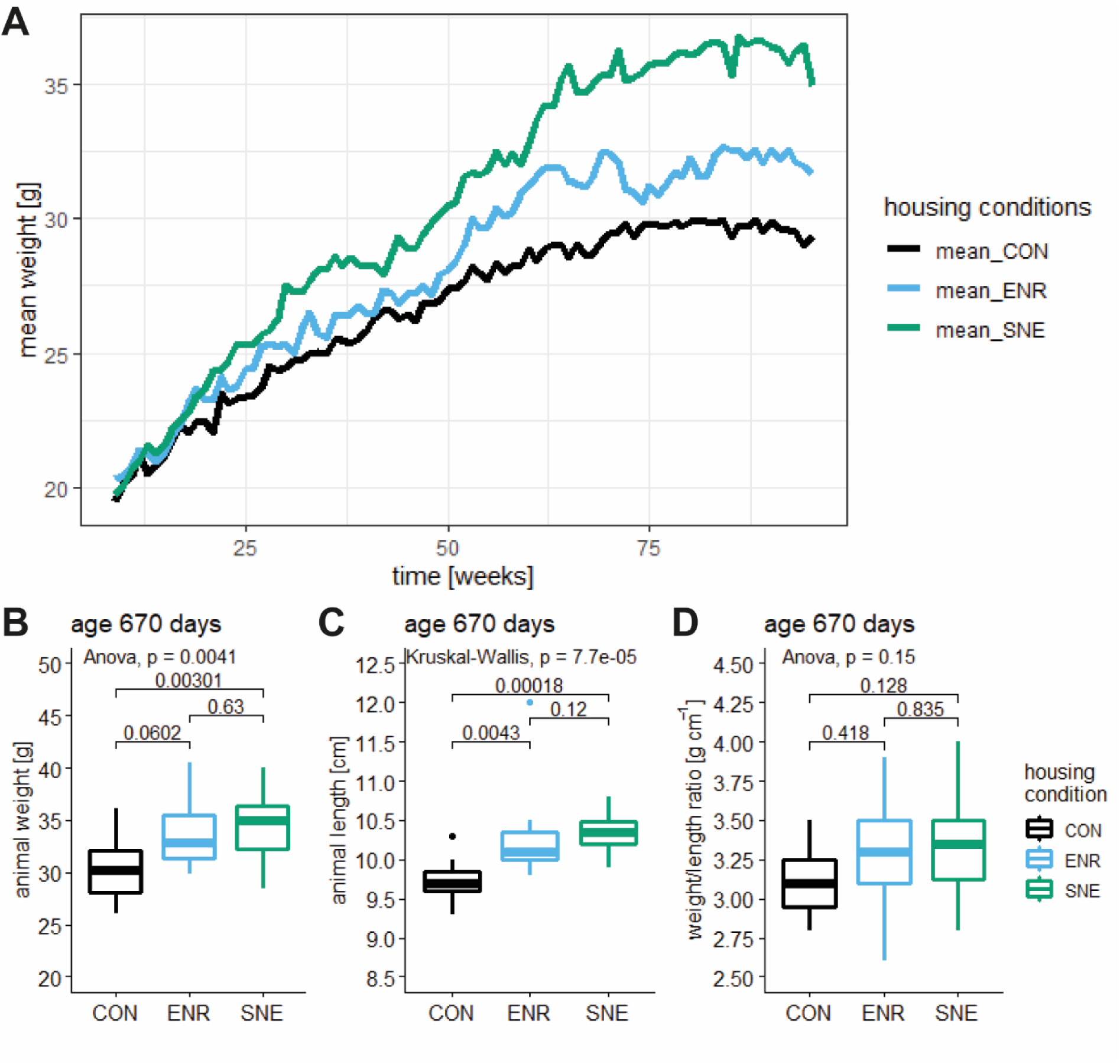
Line plot and boxplots of the physical parameters weight and body length of female C57BL/6J mice from the three different housing conditions. A – mean body weight over time with CON housing in black, ENR housing in light blue and SNE housing in green. B – mean body weight at the day of perfusion (age 670 days, same coloring scheme as A) with respective *p* values from a post hoc Tukey test. C – mean animal length at the day of perfusion (age 670 days, colors equal to A) with respective *p* values from a post hoc Wilcoxon test. D – mean ratio of animal weight to animal length at the day of perfusion (age 670 days, same coloring scheme as A) with respective *p* values from a post hoc Tukey test.

In order to be able to map the physical characteristics more accurately, the length of experimental animals was also measured and placed into relation with the animal weight. The animal length in CON housing was 9.7 ± 0.3 cm (2.8 %, 10.3 – 9.3 cm, n = 11), in ENR housing 10.1 ± 0.2 cm (2.2 %, 10.5 – 9.8 cm, n = 10) and in SNE housing 10.3 ± 0.2 cm (2.1 %, 10.8 – 9.9 cm, n = 18, Figure 1 B). Comparable to the body weight ENR and SNE animals were significantly longer than CON animals with no significant difference between them. The ratio of animal weight to animal length in CON housing resulted to 3.1 ± 0.3 g cm^-1^ (8.2 %, 3.5 – 2.8 g cm^-1^, n = 11), in ENR housing to 3.3 ± 0.4 g cm^-1^ (10.9 %, 3.9 – 2.6 g cm^-1^, n = 11) and in SNE housing to 3.3 ± 0.3 g cm^-1^ (8.7%, 4 – 2.8 g cm^-1^, n = 18, Figure 1 B). Overall, the same trend as in body weight and length was revealed, but no significant differences.

### 2.2. Adipose tissue weight

At the time of perfusion, retroperitoneal adipose tissue weight for animals in the CON housing was 0.057 ± 0.032 g (56.8 %, 0.127 – 0.030 g, n = 11), for ENR housing 0.076 ± 0.019 g (25.3 %, 0.092 – 0.037 g, n = 10) and for SNE housing 0.112 ± 0.048 g (43.1 %, 0.201 – 0.020 g, n = 18, Figure 2 A). For periovarian adipose tissue weight animals in CON housing showed 0.276 ± 0.157 g (56.9 %, 0.593 – 0.020 g, n = 11), for ENR housing 0.409 ± 0.182 g (44.5 %, 0.798 – 0.184 g, n = 11) and for SNE housing 0.506 ± 0.236 g (46.6 %, 1.053 – 0.127 g, n = 18, Figure 2 B). For both tissues SNE animals showed significant heavier adipose tissue weights than CON animals. There was no statistical difference between CON and ENR animals nor ENR and SNE animals. The weights of the adipose tissues in relation to the body weight of the mice were in the same proportion to each other. The percentage of retroperitoneal and periovarian adipose tissue increased significantly from the CON housing to the SNE housing.

**Figure 2.**
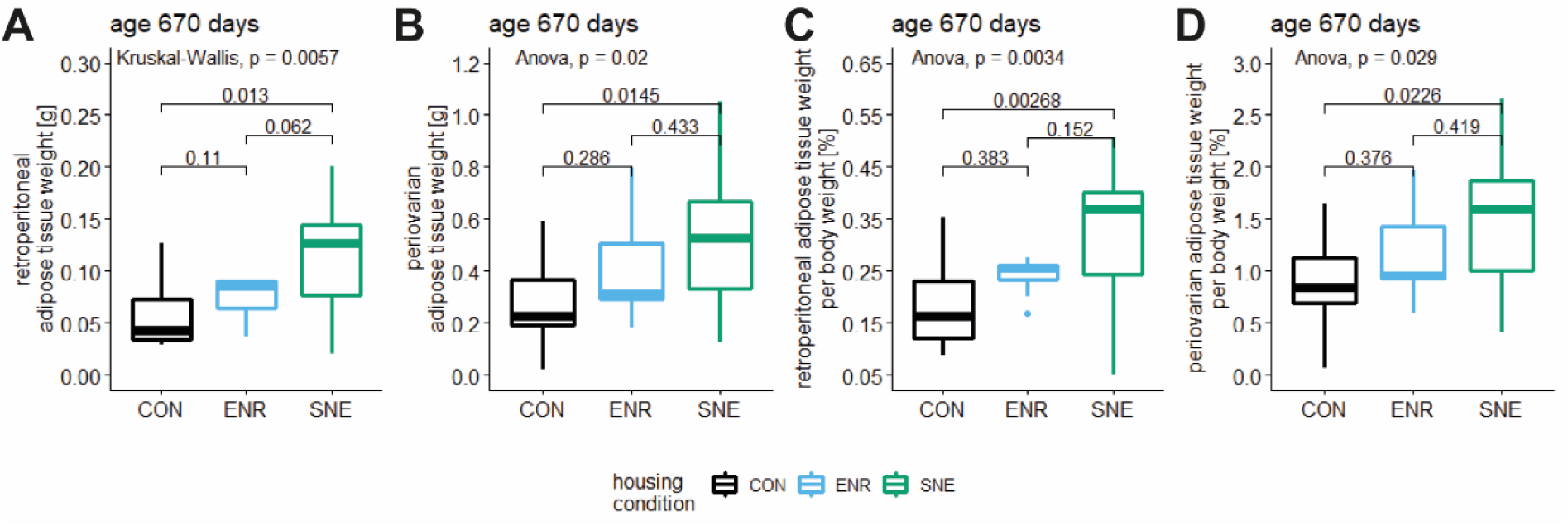
Boxplot of adipose tissue weight and adipose tissue weight relative to animal body weight of female C57BL/6J mice at the time of perfusion (age 670 days). A – retroperitoneal adipose tissue weights of animals in CON housing (black), ENR housing (light blue) and SNE housing (green) after perfusion at an age of 670 days with respective *p* values from post hoc Wilcoxon test. B – periovarian adipose tissue weights (same coloring scheme) after perfusion at an age of 670 days with the respective *p* values from post hoc Tukey test. C – retroperitoneal adipose tissue weights relative to animal body weight (same coloring scheme) after perfusion at an age of 670 days with respective *p* values from post hoc Tukey test. D – periovarian adipose tissue weights relative to animal body weight (same coloring scheme) after perfusion at an age of 670 days with the respective *p* values from post hoc Tukey test.

### 2.3. Bone density and structural properties data

To observe the change of the bone density over time, it was measured at three different ages of female C57BL/6J mice during housing in different conditions. At an age of 340 days the mice showed a bone density in CON housing of 1.51 ± 0.24 g cm^-3^ (15.6 %, 1.81 – 0.98 g cm^-3^, n = 11), in ENR housing of 1.69 ± 0.17 g cm^-3^ (16.5 %, 1.90 – 1.42 g cm^-3^, n = 12) and in SNE housing of 1.72 ± 0.16 g cm^-3^ (16.0 %, 1.92 – 1.35 g cm^-3^, n = 20, Figure 3 A). Both animals in ENR and SNE housing had a significantly higher bone density than the control animals, but showed no statistical difference between the two enriched housing conditions. At an age of 501 days the bone density values decreased to 1.44 ± 0.15 g cm^-3^ (10.6 %, 1.63 – 1.16 g cm^-3^, n = 11) in CON housing, 1.58 ± 0.19 g cm^-3^ (12.0 %, 1.87 – 1.25 g cm^-3^, n = 10) in ENR housing and 1.57 ± 0.23 g cm^-3^ (14.4 %, 2.09 – 1.36 g cm^-3^, n = 19, Figure 3 A) in SNE housing. The difference between the three housing conditions was not statistically significant anymore. Bone density showed further degression at an age of 664 days with 1.21 ± 0.13 g cm^-3^ (10.8 %, 1.44 – 1.02 g cm^-3^, n = 11) for CON animals, 1.38 ± 0.25 g cm^-3^ (17.8 %, 1.92 – 1.06 g cm^-3^, n = 11) for ENR animals and 1.35 ± 0.16 g cm^-3^ (12.2 %, 1.68 – 1.05 g cm^-3^, n = 17, Figure 3) for SNE animals. Analysis showed no significant difference although the relation between values showed a trend comparable to the first measurement at 340 days of age.

**Figure 3.**
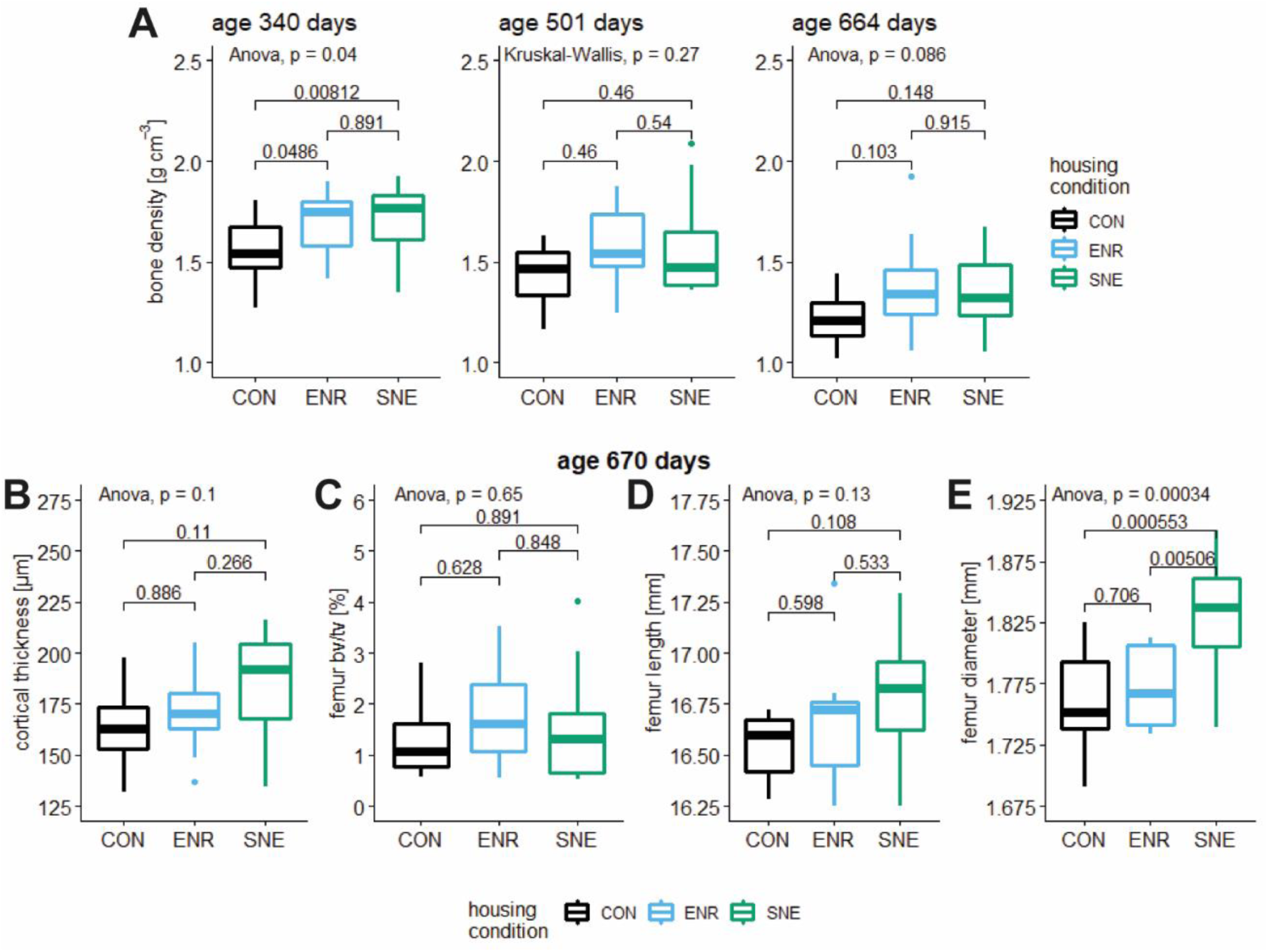
Bone density at three different ages and structural femur properties of female C57BL/6J mice in three different housing conditions. A – bone density in g cm^-3^ at the age of 340, 501 and 664 days of animals in CON housing (black), ENR housing (light blue) and SNE housing (green) with respective *p* values from post hoc Tukey (A left and right) and Wilcoxon (A middle) test. B—E – parameters were measured after perfusion at an age of 670 days in CON housing (black, ENR housing (light blue) and SNE housing (green). B – cortical thickness in µm with respective *p* values from post hoc Tukey test. C – femur bone volume per tissue volume in % with respective *p* values from post hoc Tukey test. femur midshaft outer diameter in mm with respective *p* values from post hoc Tukey test. D – femur length in mm with respective *p* values from post hoc Tukey test. E – femur midshaft outer diameter in mm with respective *p* values from post hoc Tukey test.

A linear model confirmed the significant decrease of bone density over time (F(2|120) = 27.41, p-value < 0.001, R² = 0.30). The observation of a lower bone density at a higher age of female C57BL/6J mice was mostly influenced by the housing condition as a random factor (model 1). A mixed effect model without housing condition as a random factor (model 2) was significantly less adequate to describe the data (p = 0.016, model 1 AIC = −44.35, model 2 AIC = −40.62)). However, a comparison with housing condition alone as a random factor (model 3) showed no significant difference (p = 0.24).

In addition to the bone density, detailed properties of the bone structure were determined via µCT analysis. The examination of the macroscopic compartments of the bone revealed a cortical thickness of the femur for female C57BL/6J mice in CON housing of 164.4 ± 23.2 µm (14.1 %, 197.4 – 132.1 µm, n = 9), in ENR housing 169.5 ± 19.7 µm (11.6 %, 205.3 – 136.7 µm, n = 9) and in SNE housing 185.5 ± 24.5 µm (13.2 %, 216.2 – 134.6 µm, n = 12, Figure 3 B). SNE animals showed the highest cortical thickness but the difference between the housing conditions was not significant. The ratio of trabecular bone volume to tissue volume was the lowest in CON housing with 1.32 ± 0.78 % (59.0 %, 2.81 – 0.57 %, n = 8). The highest bv/tv was measured for ENR housing with 1.78 ± 1.09 % (61.1 %, 3.52 – 0.55 %, n = 9). Animals in SNE housing showed bv/tv of 1.53 ± 1.10 % (71.8 %, 4.01 – 0.51 %, n = 18, Figure 3 C). The differences between the groups were not significant.

The femur length for the experimental animals in CON housing was 16.55 ± 0.15 mm (0.9 %, 16.72 – 16.28 mm, n = 9), was increased in ENR housing with 16.67 ± 0.32 mm (1.9 %, 17.34 – 16.25 mm, n = 9) and reached the highest value in SNE housing with 16.79 ± 0.28 mm (1.7 %, 17.29 – 16.25 mm, n = 11, Figure 3 D). The increase in length with the rising level of enrichment was not significant.

Comparable to the femur length, the femur midshaft outer diameter was also increased in ENR and SNE housing. In CON housing the diameter was 1.76 ± 0.04 mm (2.3 %, 1.83 – 1.69 mm, n = 9), in ENR housing 1.77 ± 0.03 mm (1.8 %, 1.81 – 1.73 mm, n = 9) and in SNE housing 1.83 ± 0.04 mm (2.3 %, 1.90 – 1.74 mm, n = 12, Figure 3 E). The mean diameter in SNE housing was significantly higher than the diameter in animals in CON and ENR housing.

Additional structural properties are mentioned in table (see 3.9. data summary). Respective figures are shown in supplements figure S1. In summary, no significant differences were found in the femora for cortical porosity, trabecular thickness, number and separation.

Structural bone anomalies were found in animals of every housing condition. In percentage terms, SNE housing showed the lowest number of animals with anomalies (Figure 4 D). In CON housing every individual showed at least one atypical feature (Figure 4 B **and** C). The effect of housing condition on the percentage of animals with anomalies was not significant. On average, the animals in ENR housing had 2.2 anomalies per individual animal. CON housing showed only slightly less with 1.9 anomalies per animal whereas animals in SNE housing on average showed 1.0 anomaly per individuum (Figure 4 E). Relative to the number of individuals in the housing conditions, SNE housing resulted in the fewest anomalies.

**Figure 4.**
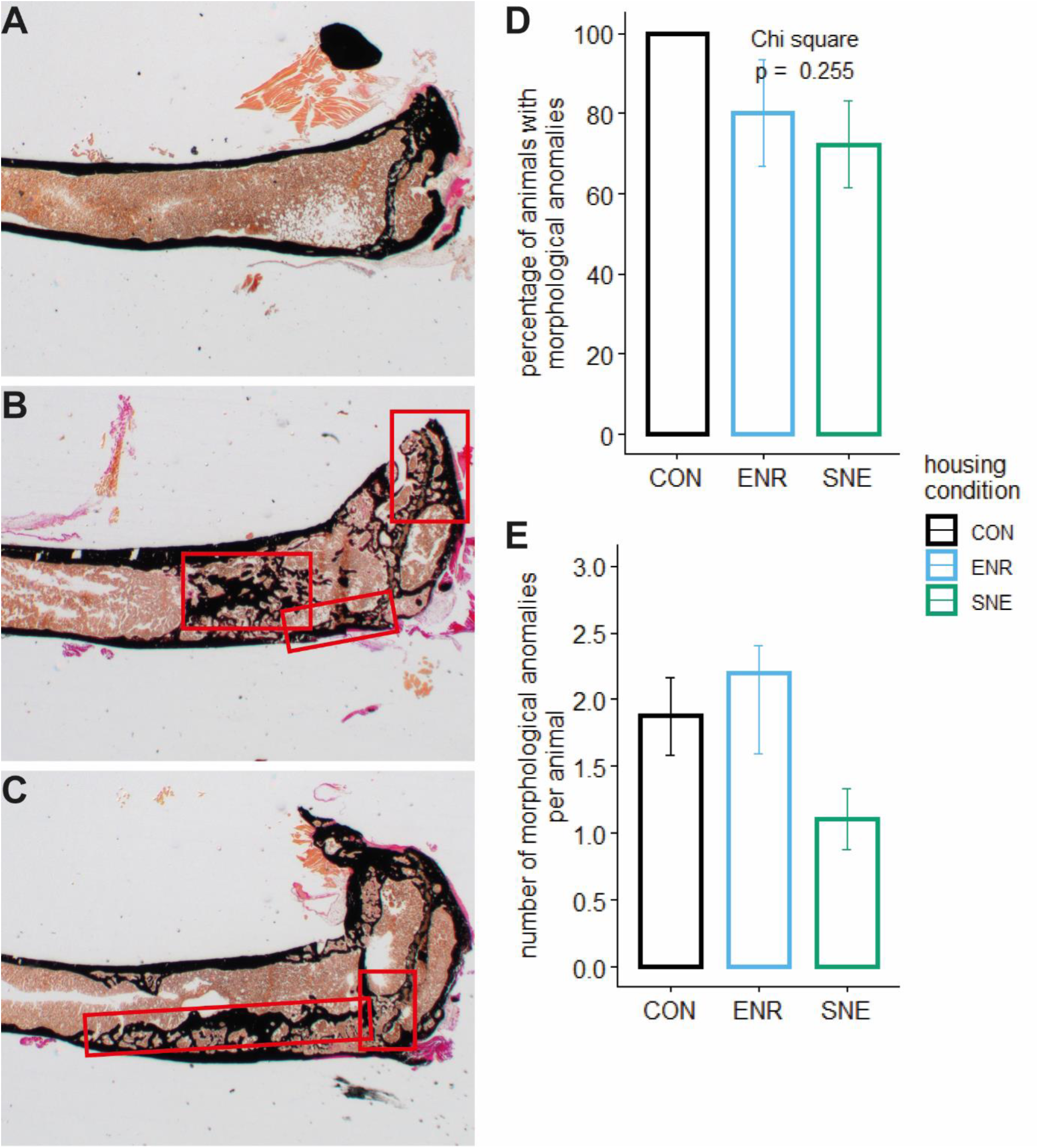
Examples of Kossa stained tibia sections (2.5×) of female C57BL/6J mice and bar plots of the occurrence of anomalies in their bone structure. A – example for no anomalies in the tibia section. B – example for trabecularized cortical bone, mineralized lesions and anomalous trabecular localization in the tibia (here: epiphysis). C – example for pseudocortical structures within the medullar cavity and the partial disruption of the growth plate. D – bar plot of the percentage of animals within each housing condition CON (black), ENR (light blue) and SNE (green), that showed morphological anomalies with the respective p value from a *χ²* test. E – bar plot of the number of morphological anomalies per animal within the three housing conditions (same coloring scheme).

### 2.4. Grip strength data

The grip strength of female C57BL/6J mice was measured at two times during their housing period. At an age of 508—510 days the mice showed a grip strength in CON housing of 2.29 ± 0.39 N (17.0 %, 3.26 – 1.78 N, n = 10), in ENR housing 2.60 ± 0.41 N (15.6 %, 3.23 – 1.99 N, n = 12) and in SNE housing 2.38 ± 0.35 N (14.5 %, 3.29 – 1.89 N, n = 19, Figure 5). Mice in ENR housing showed the highest grip strength in comparison to the animals of the other housing conditions. The difference is not significant. In the second measurement at an age of 664 days animals showed in CON housing 2.34 ± 0.43 N (18.4 %, 3.00 – 1.71 N, n = 11), in ENR housing 2.65 ± 0.26 N (9.8 %, 3.04 – 2.21 N, n = 12) and in SNE housing 2.40 ± 0.40 N (16.6 %, 3.09 –1.85 N, n = 18, Figure 5). The relationship between values has not changed and was still not significant. Also a linear model showed no significant change of the grip strength between the two measurements (F(1|79) = 0.095, p-value = 0.76, R² = −0.01).

**Figure 5.**
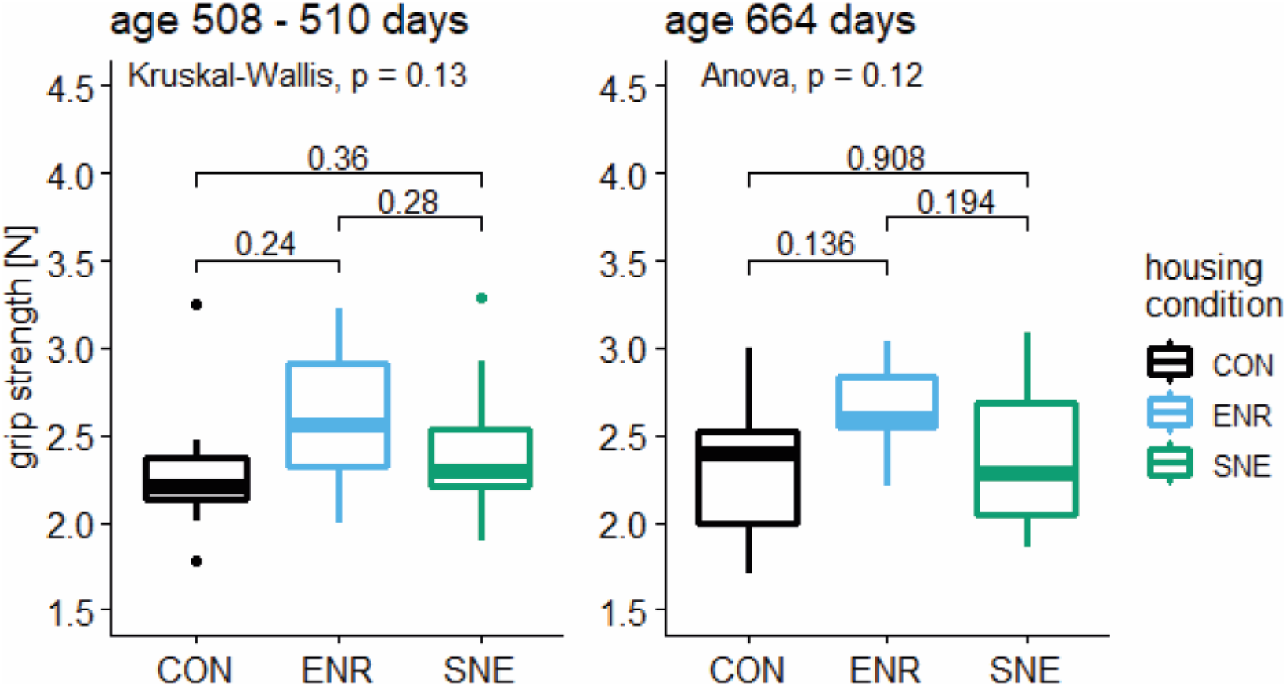
Boxplots of the grip strength of female C57BL/6J mice at two different times during housing in there different conditions. Left boxplot – grip strength at an age of 508—510 days of animals in CON housing (black), ENR housing (light blue) and SNE housing (green) with respective p values from post hoc Wilcoxon test. Right boxplot – grip strength at an age of 664 days (same coloring scheme) with respective p values from post hoc Tukey test.

### 2.5. Muscle weight data

After perfusion of the animals at an age of 670 days the muscle weight of the musculus biceps femoris was measured. For CON housing animals showed a muscle weight of 0.123 ± 0.023 g (19.0 %, 0.161 – 0.097 g, n = 11), for ENR housing 0.127 ± 0.021 g (16.5 %, 0.166 – 0.097 g, n = 11) and for SNE housing 0.120 ± 0.020 g (16.3 %, 0.161 – 0.087 g, n = 18, Figure 6). Animals in the ENR housing showed the highest muscle weight. There was no significant difference between housing conditions. Muscle weight in relation to body weight decreased significantly from the CON housing to the ENR housing to the SNE housing. However, the individual comparison between the housing conditions did not show any singificant differences.

**Figure 6.**
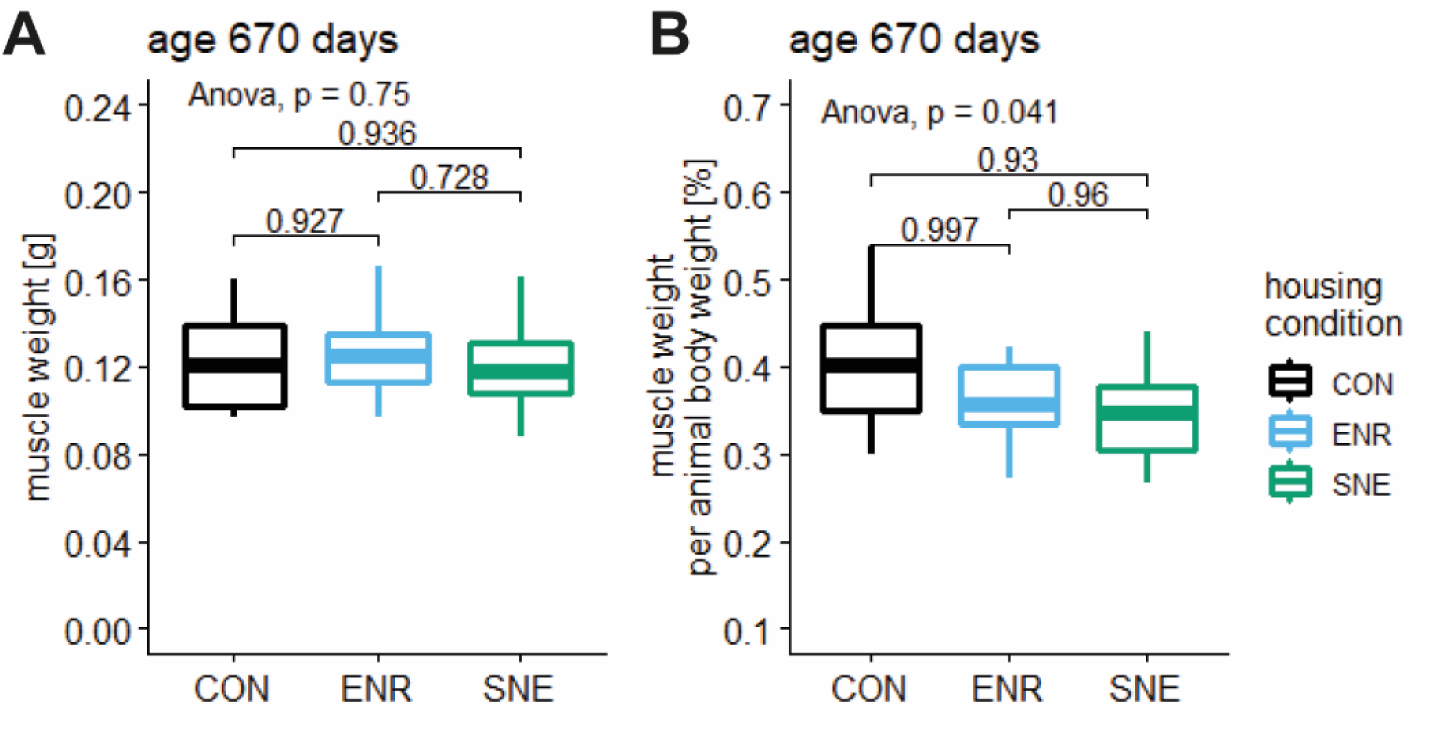
Boxplot of muscle weight in g (A) and muscle weight relative to animal body weight in % (B) of female C57BL/6J mice in three different housing conditions. (CON housing in black, ENR housing in light blue and SNE housing in green). The parameter was measured after the perfusion at the age of 670 days and is displayed with the respective *p* values from a post hoc Tukey test.

### 2.6. Bone turnover parameter data

Three musculoskeletal turnover parameters were measured in blood samples after the perfusion of female C57BL/6J mice at an age of 670 days. Myostatin is an endogenous protein that inhibits muscle growth. The myostatin concentration for animals in CON housing was 692.26 ± 132.87 pg ml^-1^ (19.2 %, 952.38 – 466.68 pg ml^-1^, n = 11), in ENR housing 714.96 ± 107.04 pg ml^-1^ (15.0 %, 796.48 – 584.37 pg ml^-1^, n = 5) and in SNE housing 692.26 ± 132.87 pg ml^-1^ (19.2 %, 952.38 – 466.68 pg ml^-1^, n = 18, Figure 7). There was no significant difference in the myostatin concentration between the three housing conditions.

**Figure 7.**
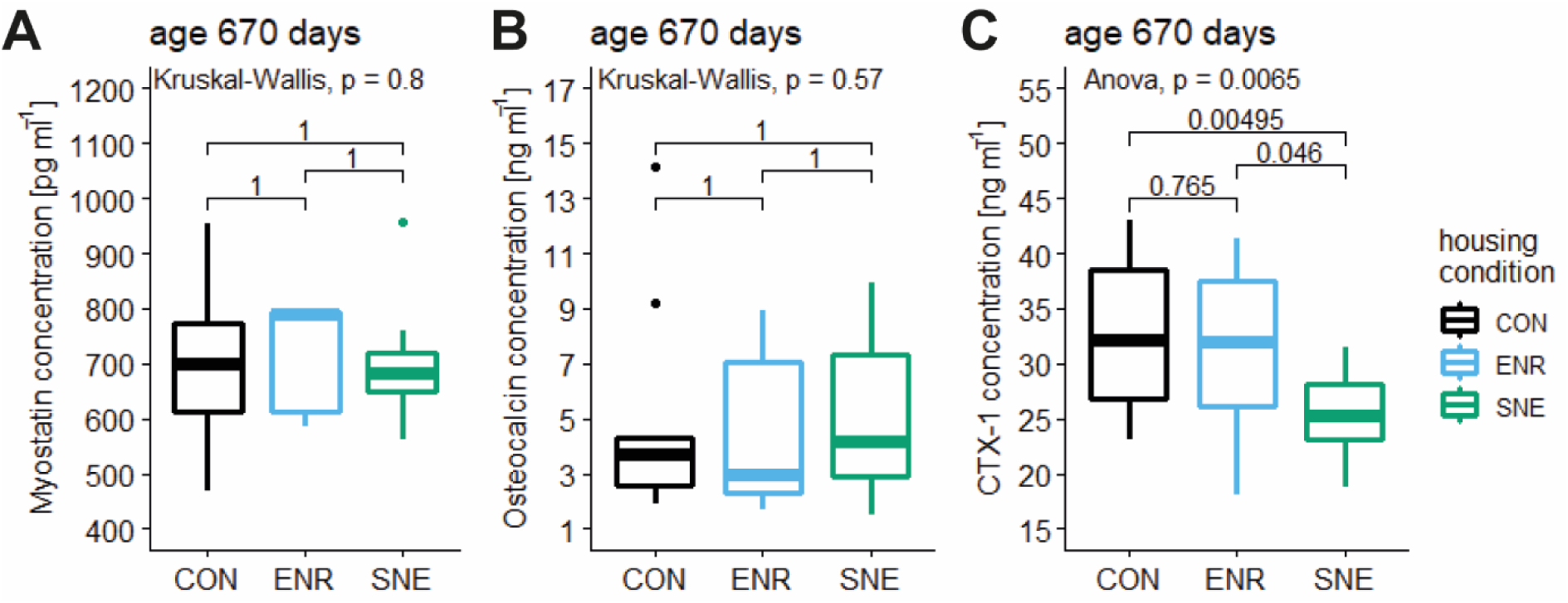
Boxplots of the blood serum concentration of three bone turnover markers myostatin (A), osteocalcin (B) and CTX-1 (C) of female C57BL/6J mice in three different housing conditions. (CON housing in black, ENR housing in light blue and SNE housing in green). The parameters were measured after the perfusion at an age of 670 days and are displayed with the respective *p* values from post hoc Wilcoxon (A and B) and Tukey (C) tests.

Osteocalcin is a component of the extracellular non-collagenous bone matrix and serves to mineralize bone during new bone formation or healing. In mice osteocalcin also stimulates insulin secretion and thus lipolysis. The osteocalcin concentration in CON housing was 4.77 ± 3.67 ng ml^-1^ (77.0 %, 14.15 – 1.90 ng ml^-1^, n = 11), lower in ENR housing with 4.48 ± 2.96 ng ml^-1^ (66.1 %, 8.95 – 1.70 ng ml^-1^, n = 9) and slightly increased in SNE housing with 5.18.26 ± 2.79 ng ml^-1^ (53.8 %, 9.91 – 1.47 ng ml^-1^, n = 17, Figure 7). Comparable to the relation in the values for the myostatin concentration, no significant difference was found.

C-terminal telopeptides are metabolic products of collagen and represent a suitable marker for bone resorption. An elevated CTX-1 level indicates a reduced bone turnover. The results for the CTX-1 concentration showed a more distinct relationship. In CON housing CTX-1 concentration was the highest with 32.97 ± 7.20 ng ml^-1^ (21.8 %, 43.08 – 23.15 ng ml^-1^, n = 10). With no significant difference to CON housing, concentration for animals in ENR housing was 31.00 ± 7.80 ng ml^-1^ (25.2 %, 41.41 – 18.14 ng ml^-1^, n = 9). SNE housing lead to a significantly lower CTX-1 concentration in comparison to CON and ENR housing with 24.77 ± 4.38 ng ml^-1^ (17.7 %, 31.49 – 14.71 ng ml^-1^, n = 18, Figure 7).

### 2.7. Resting metabolic rate

RMR data for animals in SNE housing were already published (21). Here the values were compared with those of the other housing conditions. The resting metabolic rates were measured at an age of 585—622 days. For animals in CON housing a rate of 42.5 ± 7.4 ml min^-1^ kg^-1^ (17.4 %, 50.9 – 30.6 ml min^-1^ kg^-1^, n = 10) was observed. In ENR housing oxygen consumption was at 36.9 ± 4.5 ml min^-1^ kg^-1^ (12.2 %, 47.3 – 30.2 ml min^-1^ kg^-1^, n = 10) and in SNE housing at 40.5 ± 3.4 ml min^-1^ kg^-1^ (8.4 %, 46.6 – 33.3 ml min^-1^ kg^-1^, n = 19, Figure 8). The housing condition has a significant effect on the metabolic rate of experimental animals. In ENR housing the RMR is significantly lower than in CON housing. Although also being lower than in CON housing, the decreased oxygen consumption in SNE housing is not significantly different to the rates in the other housing conditions. Animals in the CON housing showed the highest variance in the RMR.

**Figure 8.**
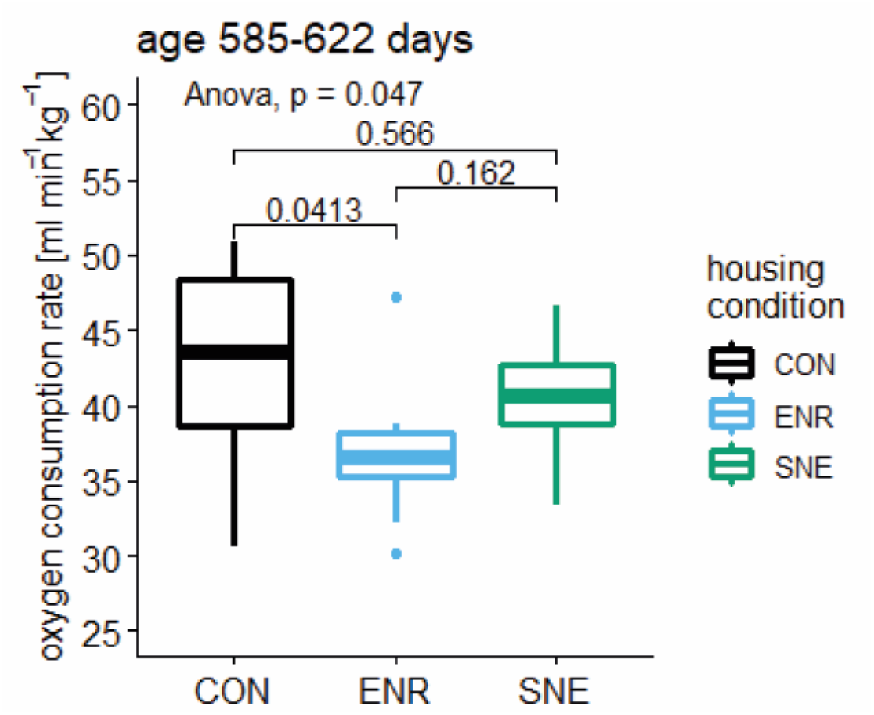
Boxplot of resting metabolic rate in mmol min-1 kg-1 for female C57BL/6J mice in three different housing conditions. (CON housing in black, ENR housing in light blue and SNE housing in green). The rates were measured at an age of 585—622 days and are displayed with the respective *p* values from a post hoc Tukey test.

### 2.8. Corticosterone and corticosterone metabolite concentration

Fecal corticosterone metabolite concentration of animals in CON housing was 2.53 ± 0.77 ng mg^-1^ (30.5 %, 3.90 – 1.26 ng mg^-1^, n = 9) and 2.73 ± 0.78 ng mg^-1^ (28.5 %, 3.85 – 1.26 ng mg^-1^, n = 11) for animals in ENR housing. Concentration was lower for animals in SNE housing with 1.94 ± 0.65 ng mg^-1^ (33.6 %, 3.66 – 0.86 ng mg^-1^, n = 19, Figure 9). There was an overall effect of housing condition with SNE mice showing significant lower fecal corticosterone metabolite concentrations than ENR animals, but not animals in CON housing. FCM concentration was not significantly different between CON and ENR housing.

**Figure 9.**
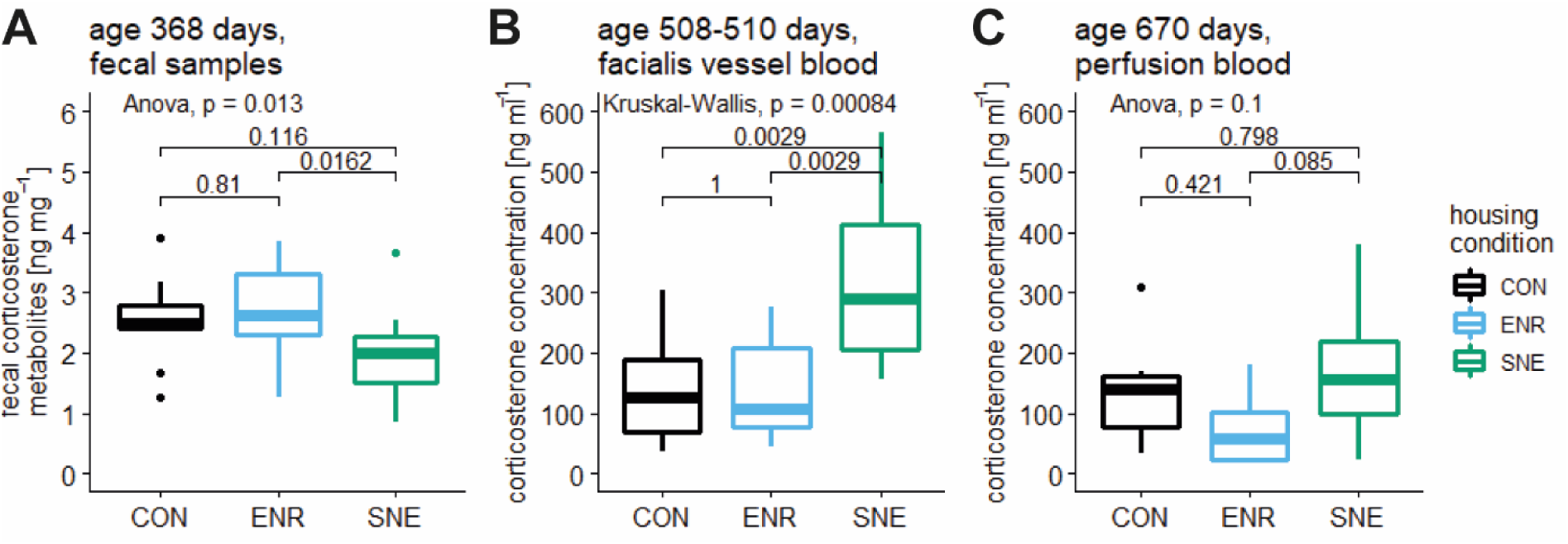
Boxplots of concentrations of fecal corticosterone metabolites (FCM) and blood corticosterone for female C57BL/6J mice in three different housing conditions. A – fecal corticosterone metabolite concentration in ng mg^-1^ in CON (black), ENR (light blue) and SNE (green) housing at an age of 368 days with respective *p* value from a post hoc Tukey test. B – facialis vessel blood corticosterone concentration in ng ml^-1^ in the housing conditions (colors equal) at an age of 508—510 days with respective *p* value from a post hoc Wilcoxon test. C – perfusion blood corticosterone concentration in ng ml^-1^ in the housing conditions (colors equal) after the perfusion with respective *p* value from a post hoc Tukey test.

Blood corticosterone concentration was measured following two different methods of blood sampling at two different time points. At 508—510 days of age, facialis vessel blood of animals in CON housing had 142.32 ± 93.33 ng ml^-1^ (65.5 %, 303.55 – 37.42 ng ml^-1^, n = 10) corticosterone. ENR housing showed 132.44 ± 81.47 ng ml^-1^ (61.5 %, 276.49 – 44.35 ng ml^-1^, n = 9). These concentration values doubled for the animals in SNE housing with 321.50 ± 139.66 ng ml^-1^ (43.4 %, 564.08 –155.03 ng ml^-1^, n = 15, Figure 9). Post hoc analysis showed that corticosterone concentration in SNE housing was significantly higher than in animals from CON and ENR housing.

After the perfusion of the animals the measurement was done with perfusion blood samples. Corticosterone concentration was 136.95 ± 90.79 ng ml^-1^ (66.3 %, 308.13 – 34.85 ng ml^-1^, n = 7) in CON housing, 78.95 ± 61.60 ng ml^-1^ (78.0 %, 179.75 – 19.26 ng ml^-1^, n = 9) in ENR housing and 163.47 ± 102.92 ng ml^-1^ (63.0 %, 379.90 – 21.75 ng ml^-1^, n = 15, Figure 9) in SNE housing. These values gave the same trend as the measurement on facialis vessel blood, but could not reach statistically significant differences.

### 2.9. Data summary and data correlation

**Table 1.**
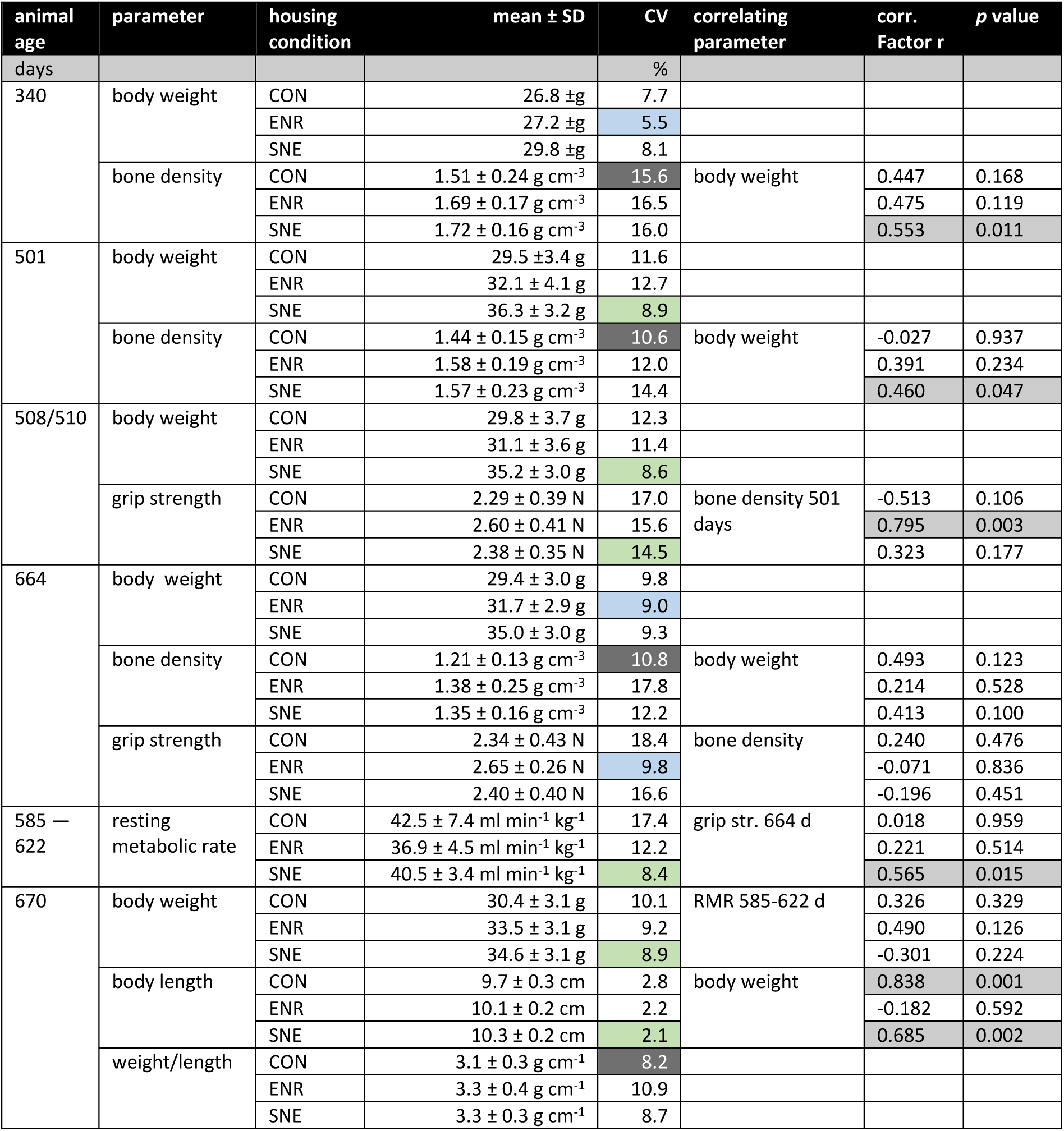
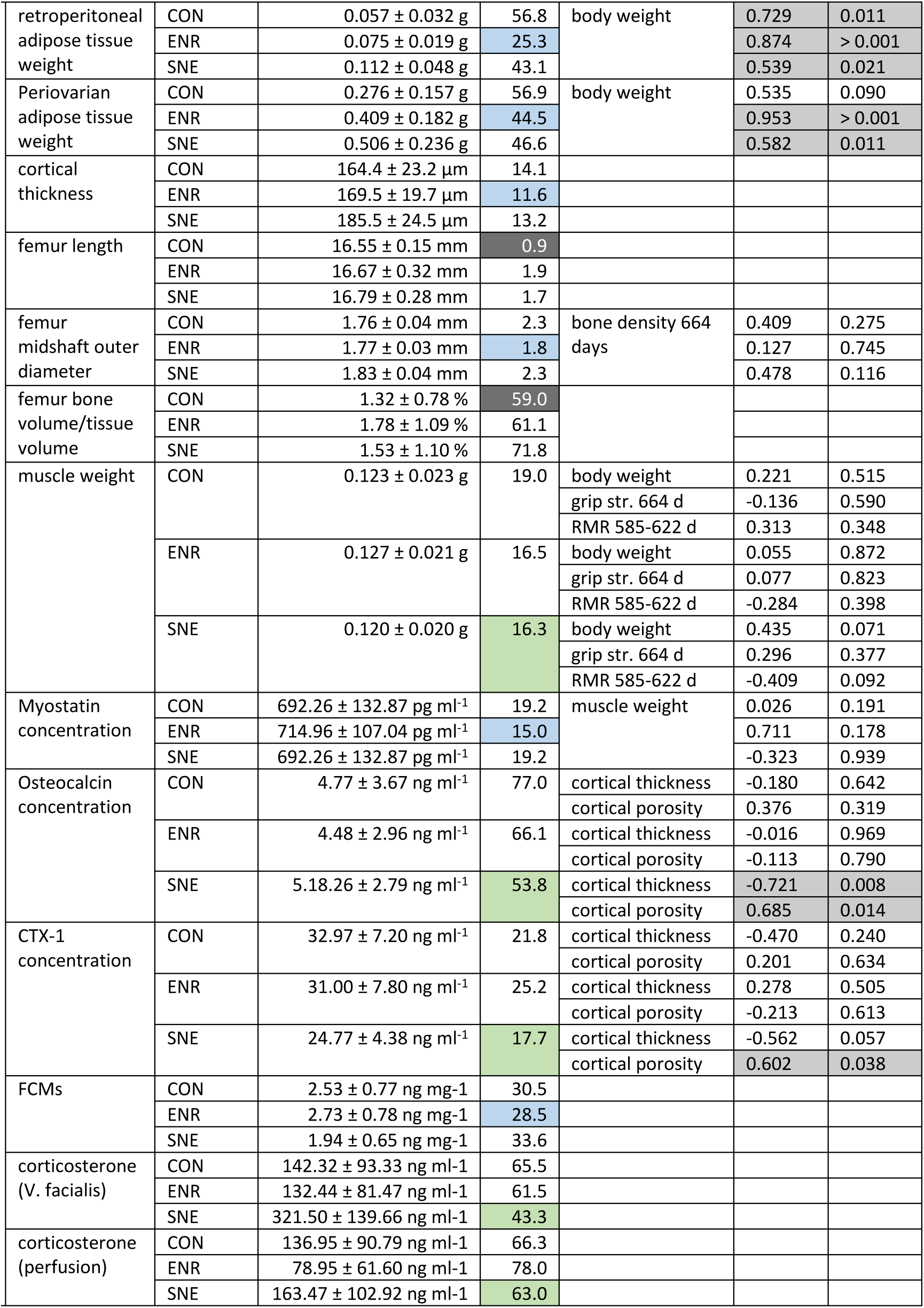
(added at the end of the manuscript): summarized results of examined parameters and correlation analysis. Shown is the age of the animals at the respective time of measurement and the value of the parameter for the animals from the three housing conditions CON, ENR and SNE housing. Values are shown as the mean with the standard deviation (SD) and the coefficient of variance (CV). The housing condition showing the lowest CV is marked in the CV column in respective to the used color scheme (CON black, ENR light blue, SNE green). If a correlation analysis with another measured value was made, it is marked as the correlating parameter and the results are shown as the correlation factor r and the respective p value. If a correlation lead to a p value lower than 0.05, the correlation is marked gray.

## 3. Discussion

Housing female C57BL/6J mice under conventional, enriched, and semi naturalistic conditions for a period of 588 days affected the examined parameters in this study in several ways. Physiological parameters body weight and body length both reached higher values with higher complexity of the housing condition. This correlated with the effect of housing condition on the examined retroperitoneal and periovarian adipose tissues. The adipose tissue weight was also higher in animals of ENR housing in comparison to CON housing and increased significantly in SNE housing. In contrast, no effect was observed in muscle weight. Comparable to the lack of differences in muscle weight, mice from different housing conditions also showed no differences in grip strength and only a weak effect on the metabolic rate was detected. Animals of the ENR housing showed the lowest oxygen demand.

The bone characteristics of the mice seem to have been influenced in part by the different housing condition. However, no uniform effect emerged. There was a clear trend of SNE housing inducing a higher bone density with a significant effect on one year old animals. This trend was also observed for the cortical thickness of the femur bones, the femur length and the femur diameter during post mortem analysis one year later. A more detailed look into bone formation and degradation parameters showed only the effect of CTX-1 being lower in blood samples of SNE animals than in CON and ENR animals indicating reduced osteoclast-mediated bone resorption. Surprisingly, all animals showed anomalies in the bone structures. However, these anomalies seemed to be reduced by the SNE housing. Finally, the stress hormones of the animals were examined according to different methods. Here, too, opposite effects were observed. In the fecal samples of the animals, the lowest corticosterone concentrations were shown in semi naturalistic environment compared to the other two housing systems. In blood samples, however, SNE animals showed the highest concentrations. The individual results are discussed in more detail below.

The conditions and experiments listed here suggest that laboratory mice achieve greater body lengths and weights with increasing complexity of housing. For body weight, this was also observed by Augustsson *et al.* in an experimental design with increasing EE (19). The difference in body weight was due to either different feed intake or different levels of exercise. Since environmental enrichment usually prevents altered feed intake (31–33) and the same type of feed was available *ad libitum* in all housing conditions, it could not be due to the feed itself but to the amount consumed and maybe to the way the feed was consumed. In cage housing, the feed was available at the feeding racks of the cage lid. In SNE, the feed was in small bowls on the bottom of the cage surface. In conventional and enriched cage housing, the animals thus reared up to get to the feed. In SNE, they eat from the floor as in nature and thereby not having to rear up. This different way of feeding might conserve energy and potentially shift energy turnover and thereby promote adipose tissue storage in the animals. However, longer distances to food sources most likely negate such an effect. The retroperitoneal adipose tissue weight did correlate with the body weight in all three housing conditions. The correlation between periovarian adipose tissue weight and body weight of the animals was not found for the CON housing condition. All in all, this indicates that feeding in enriched environments might lead to greater adipose tissue deposits and thereby higher body weight.

However, this conclusion only applies under the assumption that environmental enrichment does not affect the time spend feeding (34). In order to better understand how the increased weight in SNE and ENR conditions came about a more detailed analysis of feeding behavior is warranted. It is worth noting that none of the animals showed severe signs of obesity under any of the housing conditions.

Another reason for the physical change were more and different possibilities of movement. The ability to run longer and farther or climb more in an enriched or semi naturalistic environment suggest that more muscle is developed and therefore higher body weights are achieved (35). However, no significant difference in muscle weight was observed in relation to different body weights. One reason for this could be the advanced age of the animals at the time of the investigation. The older the animals become, the lower the muscle mass and strength of the animals (36). Regular exercise and stressing of the muscles slow down this process, but it cannot be prevented.

The lack of difference in muscle weight is consistent with the findings in the myostatin concentration of the animals. There were no significant differences in the myostatin concentration between the animals in the different housing conditions. There was also no correlation between the measured concentration of myostatin and the muscle weight of the animals in all three housing conditions. Since the muscle mass of mice is directly linked to the levels of myostatin (37, 38), an effect of housing should be recognizable on the basis of myostatin concentration. Also corresponding to the consistent muscle weight in the three housing conditions, no significant difference was detected in the grip strength of the animals. There was a comparable trend between the two measurements, in which animals of the ENR housing had the highest values of grip strength while CON and SNE had similar values. In addition, no effect of the age of the animals in relation to grip strength was observed. Muscle strength is expected to diminish with age (36, 39). However, at 558 days of age, when the grip strength was first measured the animals must already be considered old. Therefore, any effect of progressively increasing age on muscle strength could probably not have been detected.

The increasing body weight of experimental animals in this study could also have been influenced by the bone skeleton properties. This is supported by the, in some cases, significantly larger dimensions of the femora of the animals in the enriched environments. Increasing length and diameter of the femora in ENR and SNE housing were also accompanied by increasing cortical thickness. The cortical thickness and body weight did correlate in CON animals – the heavier the animals were, the thicker the femur cortical bone. It is possible, that increased opportunity for exercise in ENR and SNE housing has stimulated longitudinal growth, especially while the animals were still younger (40). Besides the mere dimensions of femora, at two different time points during the housing of the animals under different housing conditions, there was a correlation between the bone density and body weight of the animals. The higher the bone density the higher the animal’s weight. There was also an indirect correlation of cortical porosity with body weight in all three housing conditions. In CON housing even significantly before alpha correction – the heavier an animal, the less porous the femur cortical bone. However, it is likely that the underlying causality is not that mouse body weight is determined by bone density, but that higher body weight leads to higher bone density. In fact it is well established that increased body mass leads to increased bone density (41). Moreover, the correlation between body weight and bone density could not be found for CON and ENR animals.

The causes of differences in bone characteristics are multifactorial and might be influenced by movement behavior, food intake, body weight and regular forces (e.g., high jumps) experienced by the skeletal system. The increased bone density in more complex environments can on the one hand probably be explained by the larger bones in the two dimensions recorded. The SNE animals tended to have longer femora and significantly larger femora in diameter. A consistent cortical thickness was observed between the housing conditions. An equally thick bone wall with larger bones means a higher bone mass, which yields higher values in the applied bone density method. In ENR femur length did correlate with cortical thickness and did negatively correlate with cortical porosity. The longer the femur, the thicker and less porous the cortical bone. This correlation could not be shown for CON and SNE animals but might explain the effect at 340 days of age and the trends at the two later measurement times.

When looking at the parameters regarding the composition and structure of the bone skeleton, only few correlations were discovered. For ENR animals a strong correlation between bone density and grip strength was found. Animals with higher muscle loading, as it for example occurs during climbing in enriched environments, could have higher grip strength. It is known that sustained muscular loading also increases the dimensions and strength of bones (42, 43). However, contrary to our data it would be expected that in the SNE, where there was climbing at the grid and where the animals covered a lot of distance, this effect would have been confirmed.

The concentration of CTX-1 as a factor of bone resorption was significantly influenced by housing conditions. Animals living in the SNE showed less CTX-1 than the animals of CON and ENR housing. In studies on humans, exercise has been found to reduce CTX-1 even in older participants (44, 45). Therefore, it can be assumed for mice as well that the larger range of motion in the SNE reduces bone resorption. This also seems to be reflected in the frequency of bone anomalies. Although the proportion of animals exhibiting anomalies was generally high, the enriched housing had fewer animals with anomalies. These also had fewer anomalies per animal, at least in the SNE, than in the CON housing. This observation is potentially linked to the increased opportunity and necessity for movement under SNE housing conditions. It is well established that regular exercise improves skeletal metabolism by inhibiting osteoclast activity among other effects (46–48). Our results regarding CTX-1 appear to be in line with this pattern. Overall, however, only minor effects on total bone density and structural parameters were observed. It is therefore likely that other confounding factors, especially the age of the animals, mask a stronger skeletal manifestation of the observed biochemical changes.

Besides the body weight, the body length also increased in ENR and SNE housing compared to CON housing. This was examined when measuring the anesthetized animals before their perfusion at 670 days of age. A correlation was also found for body length and body weight. Except for ENR housing, all animals showed, that the longer their bodies, the higher was their body weight. This might indicate a leaner “athletic” body type under ENR conditions. The enrichment elements used in the ENR housing promotes vertical movement and the running disc should create an incentive for activity compared to the CON housing. There is indication that stereotypies, which can occur in caged mice, are compensated by increased use of running wheels (49). Therefore, increased exercise on the running disc may have favored a leaner body type. On the other hand, it remains unclear why the unequally larger exploration area in the SNE did not cause the same effect.

In general, there is a close biological relationship between resting metabolic rate and body weight. As body weight increases, greater metabolic rates are achieved within the same species. The alteration of the resting metabolic rate depends on the energy needs of all organs and body components (50). It is reasonable to assume that the possibility of more outlet and activity increases the proportional demand of muscle mass. However, in our study, muscle weight was not affected by housing conditions and the metabolic rate of mice from the SNE was not increased despite of larger cage space and possibility of movement. There was also no correlation found between body weight of the animals and their respective resting metabolic rate nor specifically between muscle weight and resting metabolic rate. To further specify the cause of the results of the present study, the activity of the animals would have to be recorded. A measure of activity in the homecage was, however, not included in our ENR and CON housing. Due to the diversity of the housing conditions, a comparable method for activity determination will not be easy.

Besides the activity of the animals, thermoregulation could be another reason for the differences in resting metabolic rate. The SNE housed twenty animals. This usually leads to a grouping of all or large groups of the mice in common nests during the resting periods. More animals within the same nest allow for less thermoregulation during resting and therefore a lower metabolic rate (51, 52). ENR housing promotes the same effect by providing additional nesting material and clearly more delineated and narrower nesting areas. On the other hand, animals in SNE housing showed no significant difference in metabolic rate compared to control animals. Lower metabolic rate due to a long-term effect of lower thermoregulation could be explained by increased stress level during calorimetry. The differences in housing conditions between SNE and single housing in a type II Makrolon measuring cage may be associated with higher stress for the SNE mice. The animals are not accustomed to a confined cage and may show increased resting metabolic rate due to increased fear.

To determine the effects of housing conditions on the stress level of the experimental animals, adrenocortical activity was measured using different samples with opposing results. Fecal samples showed a significantly lower concentration for SNE housing and equally high concentrations in CON and ENR housing. In contrast, facialis vessel blood samples showed significantly higher concentrations for SNE housing. Again, CON and ENR housing did not cause statistically relevant differences in corticosterone concentration. Results from perfusion blood samples showed no significant effect but a similar trend. SNE housing caused the highest concentration. In this measurement values for CON housing were closer to values for SNE housing than to values for ENR housing.

The fact that different results are obtained with different samples matrices is not unusual. Whereas measuring fecal corticosterone metabolites is noninvasive and of sufficient accuracy (53, 54), concentrations in fecal samples reflect a more pooled and temporally deferred sample of stress hormonal activity. FCMs are pooled due to the mixing of feces in the intestine during metabolism. The temporal delay results from the length of time the feces remain in the animals’ bodies between the measured state and the actual sampling (55, 56). In contrast, measuring corticosterone concentrations in blood always requires taking a blood sample within a few minutes. In fact, the sampling method itself introduces a potentially more stressful situation for the animals than everyday housing can cause (57–59). Although it was aimed to minimize the length of sampling, the time from picking a mouse out of the SNE until obtaining the blood sampling might have been too long to still reflect baseline corticosterone level. All in all the lack of statistical different concentrations in perfusion blood suggest that the stress level at the time of perfusion was the same for animals from all housing conditions. The results from the fecal samples on the other hand indicate that an enriched semi-natural environment reduces baseline stress levels and is in line with previous results (60). However, it should be emphasized that the different samples were also collected at different times during the housing of the mice. Thus, an effect of age as a cause for the contrasting results cannot be excluded without doubt. Given this relationship, EE seems to have more of a stress-reducing effect in this study.

Comparable to a previous analysis (21) and literature data (5), no indication for increased variability was found. To the contrary, in this study a total of 20 parameters were evaluated in 30 different measurements. In 22 measurements, lower variances of the measured values were measured for one of the two enriched housing conditions (ENR or SNE) compared to the conventional housing. In 14 cases this was true in both enriched housing conditions. This indicates that the general fear of increased variability due to improved housing conditions does not hold true.

### Limitations

Although a significant difference between the SNE housing and the other housing conditions was found in the weights of both adipose tissues examined (retroperitoneal and periovarian), these results should be viewed with caution. The variance of adipose tissue weight is strongly dependent on the execution of the section. Dissecting the adipose tissue requires skill and a clear differentiation between the target tissue and surrounding tissue. Although the preparation was always performed by the same person, methodological error cannot be ruled out.

The method to measure grip strength applied here might not have been ideal, since the steady pulling of the animals on the holding device is motorically demanding and requires training and experience of the animal as well as of the experimenter. In addition, it has to be taken into account, that during the process of pulling the animals, the motivation to hold on rather than the actual strength of the animals is measured.

One argument against the generalizability of the data collected could be the duration of the housing itself. There are a few indications in the literature that effects of EE can be reduced or weakened by long housing periods (61). Many of the parameters in our study were measured at a high age of two years. Effects of EE on the development of the animals could therefore hardly be shown. Indeed, it is likely that ageing related degenerative effects partially mask the environmental effects at this age.

This is quite well indicated by the clear and significant difference of bone density at the age of one year (generally healthy, middle-aged animals) that vanished throughout the subsequent year. Also, the high prevalence of skeletal anomalies indicates that skeletal degeneration has progressed quite far in these older mice.

On a sidenote, it can be concluded from this data that the investigated housing conditions do not prevent ageing-related bone loss.

The correlation analyses used to discuss the physiological parameters among themselves were also subject to a prior test for normal distribution. Nevertheless, these analyses involve different group sizes. In particular, the number of individuals in the SNE housing can add weight to the results. The correlations were therefore considered with caution.

### Conclusion

Overall, female C57BL/6J mice in all housing conditions exhibited strain-typical values for body weight development throughout the lifespan (62, 63). Within this range, housing them in conditions that are more natural, increased weight and length of the animals. It is worth noting, that none of the studied parameters was negatively affected by more enriched housing. All in all, bone properties appear to be slightly improved by more natural housing and age-related increased bone resorption was reduced. We confirmed previous studies, showing that the variance of the data was not increased by more natural housing conditions. This indicates that more natural housing conditions are a feasible way for housing and testing laboratory mice.

## 4. Materials and method

### 4.1. Animals

For this study 44 female C57BL/6J mice were purchased from Charles River (Charles River, Sulzfeld, Germany). Social housing of male mice in large enclosures may promote increased aggression due to territorial behavior in male animals (27). To minimize possible adverse effects of aggressive behavior, only female animals were used in this study. At arrival, animals were eight to nine weeks old. The mice were special pathogen free, were checked for their health status, weighed and then randomly assigned to one of seven groups in three different housing conditions (3 × 4 animals in conventional housing CON, 3 × 4 animals in enriched housing ENR and one group of 20 animals in a semi naturalistic environment SNE). Prior to the experiment, the groups were kept in standard Type III Makrolon cages in an open rack system. After seven days of habituation and daily handling training, animals were tagged individually with a radio frequency identification (RFID) transponder. After another two weeks of monitored recovery and handling training, animals were transferred to their respective housing conditions in a special laboratory area for animal keeping. During habituation and experimental housing animals were kept at a 12/12 h light cycle (summertime lights on 8:00 a.m.– lights off 8:00 p.m., wintertime lights on 7:00 a.m.– lights off 7:00 p.m.), at 22.0 ± 2.0 °C, and 50.0 ± 5.0 % humidity. The animals were kept in the experimental housing conditions from 82 days of age to 670 days (approx. 2 years). At different points during this time, physiological parameters were measured. Once a week, animals were weighed and handled to check for their health status. At the end of the experimental phase, the animals were put under anesthesia with a mixture of ketamine and xylazine and were transcardially perfused. Body length was measured, blood was collected, and adipose tissue, muscles, and bones were removed and weighed. Adipose tissue and muscle weights were analyzed and plotted both as actual weights and relative to animal body weight. During the study, four of the 44 animals died prior to the planned perfusion of the experimental animals due to causes unrelated to this study. The reduced animal numbers are marked in the respective parts of the results. All experiments were conducted in accordance with the applicable European and national regulations and were approved by the State Office for Health and Social Affairs Berlin (G 0069/18).

### 4.2. Transponder injection

All animals were marked individually for identification with a RFID transponder of two types (Type 1 – FDX-B transponder according to ISO 11784/85; Planet-ID, Essen, Germany/Euro I.D., Köln, Germany or Type 2 – ID 100, diameter: 2.12 mm; length: 11.5 mm, Trovan, Ltd., Douglas, UK). For analgesia the animals received the non-opioid analgesic meloxicam (0.1 mg kg^-1^, Meloxydyl, Ceva Tiergesundheit GmbH, Düsseldorf, Germany) orally 60 min before the injection. The transponder was injected subcutaneously between the shoulder blades (scapulae) under inhalation anesthesia with isoflurane according to established procedures (1.0–1.5 % in 30 % O_2_ with 70 % N_2_O). The wound was then manually closed and fixed for a few seconds to initiate natural wound closure or closed with tissue adhesive when necessary. The awakening of the animals was monitored in a separate cage.

### 4.3. Housing conditions

#### Conventional housing CON

This housing condition served as the control condition during the experiments. It meets the minimal standards for animal housing, regulated by guidelines at national and international level (i.e., directive 2010/63/EU). Similar to the standard caging during habituation CON housing consisted of a Type III Makrolon cage with a floor area of 840 cm^2^ and 153 mm height. It was filled with approx. 3 cm aspen bedding (Polar Granulate 3–5 mm, Altromin, Lage, Germany). The cage contained a red triangle plastic house, a wooden gnaw stick, a small cotton roll of nesting material and two pieces of paper towel. Mice had *ad libitum* access to tap water and food (autoclaved pellet diet, LAS QCDiet, Rod 16, LASvendi, Soest, Germany).

#### Enriched housing ENR

The enriched cage housing was set up identically to the conventional housing but was extended with different kinds of enrichment. These enrichment elements were assigned to categories regarding their prospective function and placed in the cages additionally or as an alternative to the conventional housing features. The equipment of a single cage consisted of a mouse house with a running disc, an alternative house, a wooden platform clamped between the walls of the cage, one structural element hanging from the cage lid, an interactive enrichment element and alternative nesting material in addition to the cotton roll and pieces of paper. This combination of the enrichment elements was changed every week. The only permanent element within the cage was the running disc. Elements of the other categories were combined randomly. In addition to regular food, the interactive enrichment element daily offered approx. 3.5 g millet seeds as a treatment to facilitate interaction with the interactive enrichment. The same amount of millet seeds was offered to animals in the other housing conditions (CON and SNE) by spreading it in the bedding. For a detailed description of the enrichment elements and their combination, see Hobbiesiefken et al. (2021) (3).

#### Semi naturalistic environment SNE

The SNE was set up and operated as described in Mieske et al. (2021) (21). Briefly, the SNE consists of a large mesh wired enclosure with an area of 4.6 m² spread over five different levels in different heights. On each level of the SNE there was access to water, food and shelter in form of a red triangle plastic house. The two upper levels also provided upside down Type I Makrolon cages as nesting boxes. Plexiglas tubes connected the different levels. Similar to the CON and ENR housing conditions, the SNE was also filled with 3–5 cm of aspen bedding. Every level provided two wooden gnawing sticks, two cotton rolls and two pieces of cellulose paper as nesting material. The two nesting boxes also provided two cotton rolls and two pieces of paper. For additional enrichment, the animals had access to Plexiglas tubes as structural enrichment elements and a small selection of self-designed toys of different shape and color.

### 4.4. Bone density and structural properties

X-ray images for determination of bone density were obtained on a Bruker InVivo Xtreme II (Bruker, Billerica, MA USA). The animals were picked in a randomized order, were anesthetized with isoflurane (1.0–1.5 % in 30 % O_2_ with 70 % N_2_O) and placed on the platform for X-ray acquisition. Anesthesia was maintained during the whole procedure. Eyes of the animals were protected with dexpanthenol creme. The awakening of the animals was monitored before the animals were placed back into the home cage. On the x-ray images a region of interest (ROI) was selected on the right femur of the animals. Bone density in g cm^-3^ was then determined by the Bruker Molecular Imaging Software. Bone density was measured three times during the housing of the animals at 340 days of age, 501 days and 664 days.

After perfusion of the animals, the leg bones of the animals were dissected. The samples were fixed for 24 h in paraformaldehyde (4 % PFA), washed three times with phosphate-buffered saline (PBS) and afterwards stored in 30 % sucrose solution. Length and diameter of the right femur and characteristics of the cortical and trabecular bone were analyzed using x-ray micro-computed tomography (µCT). The μCT scanning and analysis were performed as described by Zhao et al., 2021 (64). Briefly, the right femur of each mouse was fixed and placed into a radiotranslucent sample holder. Samples were scanned and analyzed with a voxel resolution of 10 μm using a μCT 40 desktop cone-beam microCT (Scanco Medical, Switzerland) according to standard guidelines (65). Trabecular bone was analyzed in the distal metaphysis in a volume situated 2500–500 μm proximal to the distal growth plate. Cortical bone was analyzed in a 1000 μm long volume situated in the middle of the diaphysis. Cortical bone evaluation was performed with a threshold of 300, whereas for trabecular bone, a threshold of 250 was used. The length of the femora was determined by the number of slices containing the bone.

For histology, tibiae were embedded in Poly(methyl methacrylate)(PMMA) and sectioned at 4 µm thickness in the sagittal plane. Sections were stained by the von Kossa/van Gieson or Toluidine blue staining procedure (66). Structural anomalies in the tibia bones were characterized by microscopic inspection and their occurrence was counted. For biomechanical testing, a three-point bending test was performed on dissected femora using a Z2.5/TN1S universal testing machine and testXpert software (both Zwick Roell, Germany) as described previously (67).

### 4.5. Grip strength

Animals were tested separately and in a randomized order. Grip strength was measured with a computerized grip strength meter (TSE Systems GmbH, Bad Homburg, Germany). The apparatus consisted of a T-shaped metal bar connected to a force transducer. To measure the grip strength in the hind paws of the mice, the mice were carefully held at the base of the tail and guided towards the metal bar with their hind paws. Their front paws were placed on a wire mesh cylinder to prevent the mice from grasping the bar with their front paws. The animal was then gently pulled backwards until the grip was lost. The peak force applied to by the hind legs was recorded in ponds (p) and converted to Newton (N). This measurement was done three times per animals on one day and the mean peak value was recorded. After the procedure, animal ware placed back into their home cage. The grip strength was measured two times during the housing of the animals at the ages of 508-510 days and 664 days.

### 4.6. Bone and muscle turnover markers

The blood serum concentration of the three following bone and muscle turnover parameters were analyzed with enzyme-linked immunosorbent assays (ELISA). All used ELISA kits were performed according to the manufacturer’s instructions.

*C-terminal telopeptides (CTX-1) –* Serum CTX-1 concentration was detected with the RatLaps™ (CTX-1) ELISA kit (competitive ELISA) (Immunodiagnostic Systems Holdings Ltd., Boldon, UK).

*Osteocalcin –* Osteocalcin concentration in the blood serum was detected with the Mouse Osteocalcin (OC) ELISA kit (competetive ELISA) (MBS275134, MyBioSource, Inc., San Diego, CA USA). The serum was diluted 1:10 before analysis.

*Myostatin –* Serum myostatin concentration was analyzed with the Mouse Myostatin ELISA kit (quantitative sandwich ELISA) (MBS166373, MyBioSource, Inc., San Diego, CA USA).

### 4.7. Resting metabolic rate

The principle of indirect calorimetry (TSE phenomaster, TSE Systems GmbH, Bad Homburg, Germany) was used to evaluate the metabolic rate. The calorimetry system measures differences in the composition of air passed individually through four measurement cages and an empty reference cage. The system was situated at a separate room at a 12/12 h light cycle, 22.0 ± 2.0 °C, and and 50.0 ± 5.0 % humidity. Animals were tested at 584–594 days of age in a randomized order. Following habituation to the experimental room (12h), mice were weighed and placed individually in measurement cages equipped with bedding, shelter and nesting material. Food and water were accessible *ad libitum* during the entire measurement and were weighed before and after the experiment. Measurement cages and the reference cage were perfused with air. In the measurement cage oxygen was lowered and carbon dioxide was increased by the respiration of the animals during the measuring period (12 h light period). After flowing through both cages, the composition of air was compared between measurement cage and reference cage. By calculating the difference between air compositions, the metabolic rate of the examined animal was assessed. After the measurement the mice, food, and water were weighed and the animals were placed back into their home cage. The resting metabolic rate (RMR) was measured as oxygen consumption rate *V̇_O_2__* during the resting phases of the animals. To separate resting phases from active phases, the cumulative frequency percentage was plotted against the measured *V̇_O_2__*. With a segmented linear regression, the threshold between *V̇_O_2__* of the resting phase and the active phase could be calculated. Data below the threshold was used to determine the RMR ((68); R package ’segmented’).

### 4.8. Corticosterone and corticosterone metabolite concentration

The concentration of corticosterone or corticosterone metabolites was measured two times during the housing of experimental animals and one time after the perfusion of the animals. The first measurement was done at an age of 368 days. At 8:00 to 10:00 am animals were individually placed in a random order in Type II Makrolon cages. The cages were just equipped with flatly spread paper towels. After a minimum period of 20 min and maximum of 30 min animals were placed back into their home cage. The fecal boli that the animals had deposited in isolation were collected and used for analyzing corticosterone metabolites (fecal corticosterone metabolites - FCM) as described before (53, 55).

At an age of 508–510 days on three consecutive days animals were individually fixated and blood was taken from the *Vena facialis* blood vessel after puncture with a lancet needle. For the CON and ENR animals, blood sampling was performed immediately after the animals were removed from the cage (within 1 minute). The SNE animals were first removed from the large enclosure and held collectively in a type 4 cage for a short time (30–45 min). The blood samples were collected in 0.2 ml reaction tubes and stored at −80 °C for further analysis. Serum corticosterone concentration was determined with a DRG Corticosterone ELISA (EIA-4164, DRG International Inc., Springfield, NJ USA). The third measurement was done with the blood samples collected after the perfusion of the animals at 670 days of age. Concentration of corticosterone was determined as described before.

### 4.9. Statistical analysis

Unless described otherwise, all measured data is presented as mean ± standard deviation. In addition, the coefficient of variation (CV), the maximum value, the minimum value and the number of measured animals is given (*x̅* ± SD (CV, max – min, n)).

Analysis and illustration of data was done with the software environment R (v 3.6.3, R Foundation for Statistical Computing, Vienna, Austria, R Studio v 1.2.1335, RStudio, Inc., Boston, MA, USA). When preparing the data for statistical analysis, they were first examined for normal distribution (‘shapiro.test()’ function) within the different groups (housing conditions). Possible outliers were identified using the ‘boxplot.stats()$out’ function and excluded for the presentation and statistical analysis of the respective data set. If normal distribution was given, an ANOVA with a Tukey post hoc analysis was used to compare the data between the housing conditions (‘aov()’ and ‘tukey_hsd()’ function). If data were not distributed normally, the Kruskal-Wallis test with a Wilcoxon post hoc analysis was applied (‘kruskal.test()’ and ‘compare_means(method = “wilcox.test”)’ function in ‘ggpubr’ package). Unless described otherwise, boxplot figures show the adjusted *p*-value after Bonferroni correction.

Continuous data were analyzed using linear models (‘lm()’ function). Related predictors were added as mixed effects to the regression models (package ‘lme4’ (69), ‘lmer()’ function). Subsequent statistical comparison of different models (‘anova()’ function) identified the factors affecting the continuous data.

Possible statistical differences in discrete data were analyzed using the χ² test (chi square test). This was performed by using the ‘chisq.test()’ function.

## Declarations

### Ethics approval and consent to participate

All experiments were approved by the Berlin state authority, Landesamt für Gesundheit und Soziales, under license No. G 0069/18 and were in accordance with the German Animal Protection Law (TierSchG, TierSchVersV).

### Consent for publication

Not applicable

### Availability of data and materials

The datasets generated and/or analysed during the current study are available in the RefinementReferenceCenter/musculoskel2022_mieskep_available_data repository, https://github.com/RefinementReferenceCenter/musculoskel2022_mieskep_available_data

### Authors’ contributions

Conceptualization, P.M., U.H., K.D. and L.L.; methodology, P.M., U.H., K.D. and L.L.; formal analysis, P.M., J.S., J.P., L.B., T.Y., R.P.; data curation, P.M.; writing—original draft preparation, P.M.; writing— review and editing, P.M., U.H., J.S., J.P., L.B., T.Y., R.P., K.D. and L.L.; visualization, P.M.; supervision, K.D. and L.L.; project administration, K.D. and L.L. All authors have read and agreed to the published version of the manuscript.

## Acknowledgements

Large parts of the SNE and the RFID tracking software were developed during L.L.’s postdoctoral period in the laboratory of N. Sachser, who generously provided all the material. The authors thank the animal caretakers, especially Carola Schwarck and Lisa Gordijenko, for their support in the animal husbandry. Special thanks and appreciation to Prof. Dr. Dieter Felsenberg for his contribution to the overall concept of the study. We thank Olga Winter, Andrea Thieke, Annette Jung and Edith Klobetz-Rassam for excellent technical assistance.

## Supplements

Additional data to – 3.3 Bone density and structural properties data is provided in Figure S1 and Table S1

**Figure S1.**
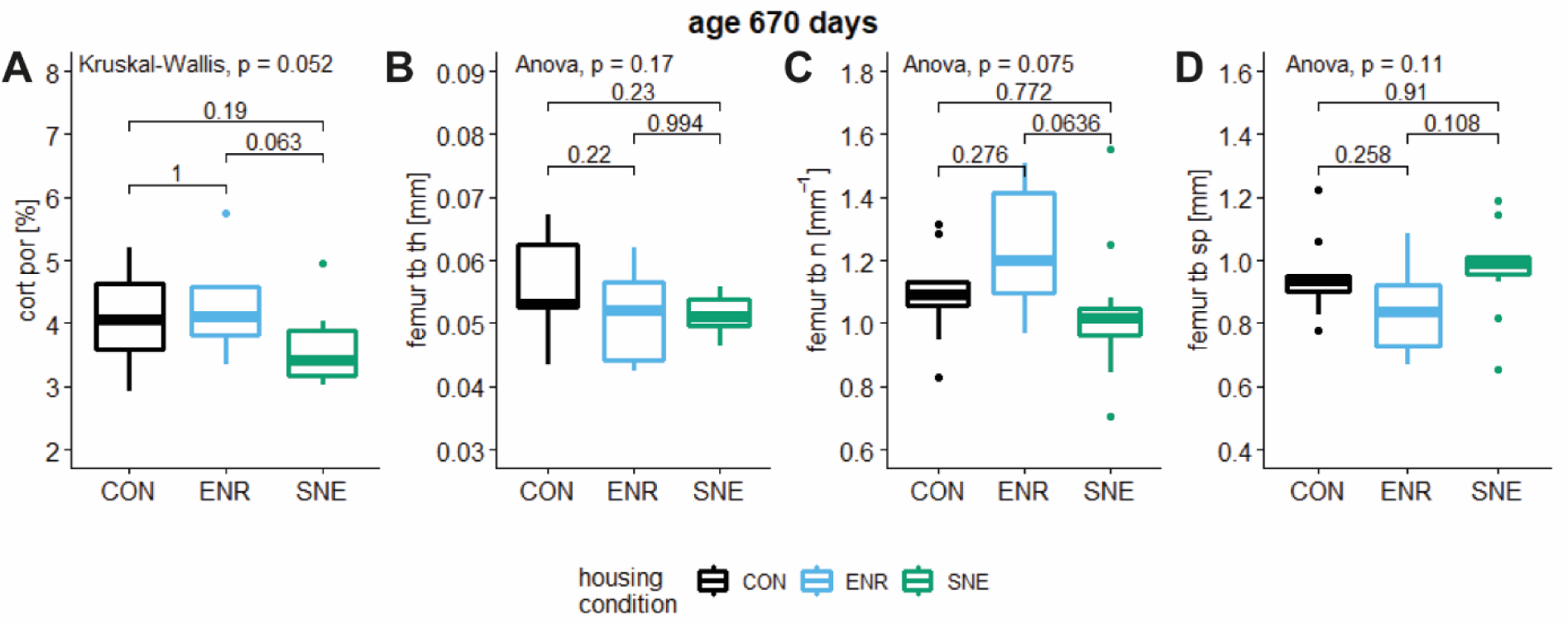
Structural femur properties of female C57BL/6J mice in three different housing conditions. A – cortical porosity in % with respective *p* values from post hoc Wilcoxon test. B – femur trabecular thickness in mm with respective *p* values from post hoc Tukey test. C – number of femur trabecular bones in mm^-1^ with respective *p* values from post hoc Tukey test. D – separation between femur trabecular bones in mm with respective *p* values from post hoc Tukey test.

**Table S1.** **Summarized results of examined parameters** in addition to 3.3 Bone density and structural properties data and Table 1. Shown is the age of the animals at the respective time of measurement and the value of the parameter for the animals from the three housing conditions CON, ENR and SNE housing. Values are shown as the mean with the standard deviation (SD) and the coefficient of variance (CV). The housing condition showing the lowest CV is marked in the CV column in respective to the used color scheme (CON black, ENR light blue, SNE green).

| animal age | parameter | housing condition | mean ± SD (max-min, n) | CV |
| --- | --- | --- | --- | --- |
| days |  |  |  | % |
| 670 | cortical porosity<br>(*Figure S1 A*) | CON | 4.1 ± 0.7 % (5.2 – 2.9 %, n = 9) | 17.6 |
| | | ENR | 4.2 ± 0.7 % (5.8 – 3.3 %, n = 9) | 16.7 |
| | | SNE | 3.6 ± 0.6 % (5.0 – 3.0 %, n = 12) | 15.5 |
| | femur trabecular thickness<br>(*Figure S1 B*) | CON | 0.056 ± 0.007 mm (0.067 – 0.044 mm, n = 9) | 13.2 |
| | | ENR | 0.051 ± 0.007 mm (0.062 – 0.042 mm, n = 9) | 14.3 |
| | | SNE | 0.051 ± 0.003 mm (0.056 – 0.046 mm, n = 11) | 6.4 |
| | femur trabecular number<br>(*Figure S1 C*) | CON | 1.09 ± 0.15 mm<sup>-1</sup> (1.31 – 0.83 mm<sup>-1</sup>, n = 9) | 13.8 |
| | | ENR | 1.23 ± 0.20 mm<sup>-1</sup> (1.51 – 0.97 mm<sup>-1</sup>, n = 9) | 16.0 |
| | | SNE | 1.03 ± 0.21 mm<sup>-1</sup> (1.55 – 0.71 mm<sup>-1</sup>, n = 12) | 20.3 |
| | femur trabecular separation<br>(*Figure S1 D*) | CON | 0.948 ± 0.131 mm (1.224 – 0.777 mm, n = 9) | 13.8 |
| | | ENR | 0.843 ± 0.140 mm ( 1.085 – 0.668 mm, n = 9) | 16.6 |
| | | SNE | 0.974 ± 0.144 mm (1.187 – 0.654 mm, n = 11) | 14.8 |

